# Predicting early bactericidal activity of tuberculosis drug combinations using a translational pharmacokinetic-pharmacodynamic modeling approach

**DOI:** 10.64898/2026.01.29.702705

**Authors:** Niurys de Castro Suarez, Eric L. Nuermberger, Jacqueline Ernest, Rada Savic

## Abstract

Phase IIa pulmonary tuberculosis (TB) trials typically assess the early bactericidal activity (EBA) of monotherapy for over 14 days. However, few studies have evaluated drug combinations, even though optimal monotherapy doses may not directly translate to combinations. Translational pharmacokinetic-pharmacodynamic (PK-PD) modeling has shown promise in predicting human treatment responses based on preclinical monotherapy data; however, its application in drug combinations remains limited. This study aimed to extend and validate our previously developed translational monotherapy PK-PD modeling platform to predict the EBA of two-drug combinations. Interactions between bedaquiline, pretomanid, linezolid, and pyrazinamide were characterized using two modeling approaches: the empirical SUPER method and the mechanistic General Pharmacodynamic Interaction model. Both approaches were independently linked to our translational platform and validated using mouse data and Phase IIa clinical results from the NC-001 study. Both modeling methods identified consistent interaction patterns, including antagonistic interactions when bedaquiline was combined with either pretomanid or linezolid. Pyrazinamide has emerged as the most effective companion for both bedaquiline and pretomanid. Our platform reasonably predicted 14-day clinical sputum colony-forming unit counts for multiple two-drug combinations, with most observations falling within the 95% prediction intervals, supporting its use in accelerating regimen development. Our study demonstrated that the translational PK–PD platform reliably predicts both short- and long-term outcomes for combinations, regardless of the interaction model. This supports its application across drug development stages to inform dose selection and effective companion drugs for anti-TB therapies.

## Introduction

Tuberculosis (TB), caused by *Mycobacterium tuberculosis* (*Mtb*), is one of the principal causes of morbidity and mortality worldwide. According to the World Health Organization (WHO), approximately 10.7 million people were affected by TB in 2024, including about 8.3 million new cases (1). Although progress has been made in diagnosis and treatment, TB remains difficult to control globally.

Standard TB treatment is based on combination therapy that targets various bacterial subpopulations and prevents the emergence of drug resistance (2). For example, WHO guidelines recommend that newly diagnosed drug-susceptible pulmonary TB be treated with an initial two-month intensive phase using isoniazid, rifampicin, pyrazinamide, and ethambutol, followed by a four-month continuation phase with isoniazid and rifampicin (3). New drugs and combination regimens are needed to shorten treatment duration, improve patient adherence, and reduce relapse rates (1). Preclinical efficacy models, particularly murine models, are essential for developing new combination regimens because they recapitulate infection and treatment responses. Although no murine model fully reproduces all relevant aspects of the disease in humans, these models play a key role in predicting human doses and extrapolating antibiotic responses, thereby guiding the development of new therapies (4, 5).

Pharmacokinetic-pharmacodynamic (PK-PD) modeling has become an essential tool for optimizing the development of therapeutic combinations because it allows robust characterization of drug interactions and more accurate prediction of human responses. However, one of the main challenges is determining which modeling approach best describes drug–drug interactions and the relative contribution of each drug within a combination. This step is crucial for identifying effective backbone regimens, particularly in TB therapy, where treatments often involve three or more drugs (6, 7). The development of advanced quantitative approaches, such as translational modeling, helps address this challenge by integrating preclinical and clinical data. By improving the extrapolation of preclinical findings, translational modeling supports dose optimization and helps reduce the need for lengthy and costly clinical trials (8, 9).

This approach enhances the efficiency of new drug development by ensuring that only the most promising combinations progress to advanced clinical studies, thereby fostering a more rational and ethical process in the fight against TB. A PK-PD platform capable of predicting Phase IIa outcomes based on preclinical efficacy in mice for drugs evaluated as monotherapy has been developed and validated in our laboratory (10). This model, integrating bacterial dynamics with exposure-response relationships in mice and human PK profiles, has demonstrated a high predictive performance for ten anti-TB drugs. Based on these results, we expanded its application to drug combinations by incorporating new model components such as PD interaction analysis.

As a first proof of concept, we previously applied this extended platform to the novel bedaquiline, pretomanid, moxifloxacin, and pyrazinamide (BPaMZ) combination (11), which has strong bactericidal activity and treatment-shortening potential needed to prevent relapse in a murine TB model (12). This study demonstrated that integrating the empirical SUPER approach, a method developed by Muliaditan *et al*. (7) to quantify pharmacodynamic interactions, enabled the use of the translational platform to successfully predict Phase IIb and III outcomes for a specific combination, representing a critical first step in adapting the model to combination therapies (11).

Using this prior work as a starting point, the present work aims to inform the generalization of this strategy by systematically evaluating two mathematical models for integrating combination drug PK-PD interactions into the translational PK-PD platform: the empirical SUPER approach (7) and the General Pharmacodynamic Interaction (GPDI) approach (13) (Fig. 1). Each approach was independently assessed to determine its ability to predict the early bactericidal activity (EBA) of five two-drug combinations based on murine data. While the current study focused exclusively on the prediction of Phase IIa outcomes, this evaluation represents an essential step toward establishing a general methodology to support dose and regimen selection using preclinical data.

**Figure 1.**
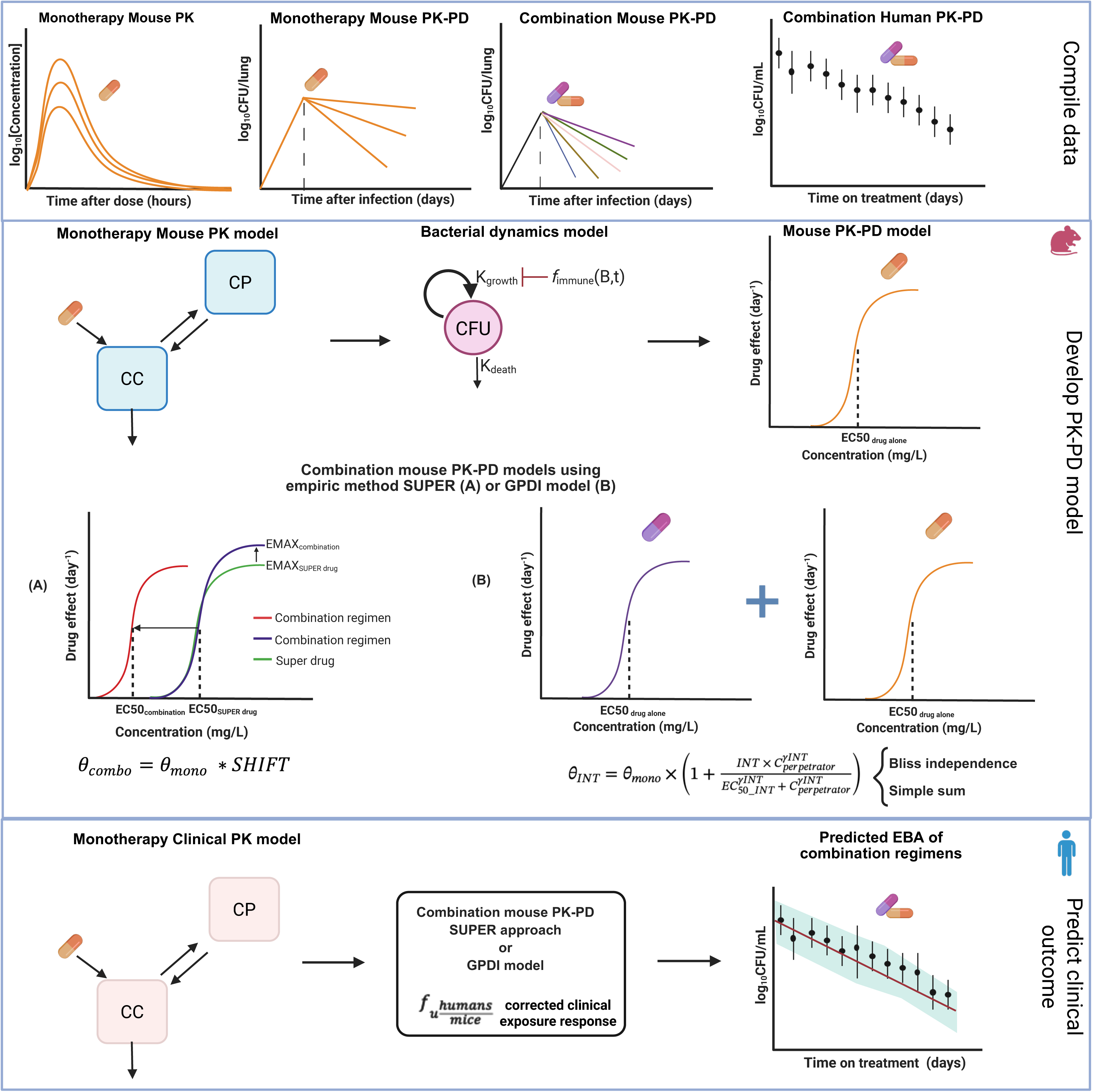
Main components of the modeling workflow: (i) Compile data and tools from both preclinical and clinical studies; (ii) Build combination mouse PK-PD models using SUPER and GPDI models; (iii) Simulate clinical outcomes using clinical PK models linked to translated exposure-response relationship. Mouse EC50 is corrected using the ratio of drug unbound fraction in humans to that in mice. A description of the equations is provided in the supplementary materials.

The SUPER method estimates PD interactions based solely on exposure-response data without requiring predefined mechanistic assumptions. It is particularly useful when experimental data are available, and a practical and rapid approach is needed to understand these interactions. In contrast, the GPDI approach provides a mechanistic modeling framework that quantifies how the concentration of one drug (the perpetrator) alters parameters such as potency (EC_50_) and maximum effect (E_max_) of another drug (the victim), offering deeper insights into the biological basis and directional nature of drug interactions. (14, 15).

## Results

### Database of two-drug combinations in mice and humans

To build the translational PK-PD model for predicting change in log₁₀ colony-forming unit (CFU) in sputum over 14 days in a Phase IIa trial, preclinical PK data (Table 1) and efficacy data in a BALB/c mouse infection model (CFU/lung, Table 2) were collected, along with population PK models and PD data in humans. The PK and efficacy of bedaquiline (B), pretomanid (Pa), linezolid (L), and pyrazinamide (Z) as monotherapy were described using previously developed mouse PK-PD models (11). The efficacy data for drug combinations in mice were gathered from experiments conducted at Johns Hopkins University (JHU) for BPa, BL, PaL, BZ, and PaZ combinations (Fig. 2A, Table 2). In total, 580 observations in mice were used to assess the reduction in CFU/lung following the administration of the therapeutic combinations.

**Figure 2.**
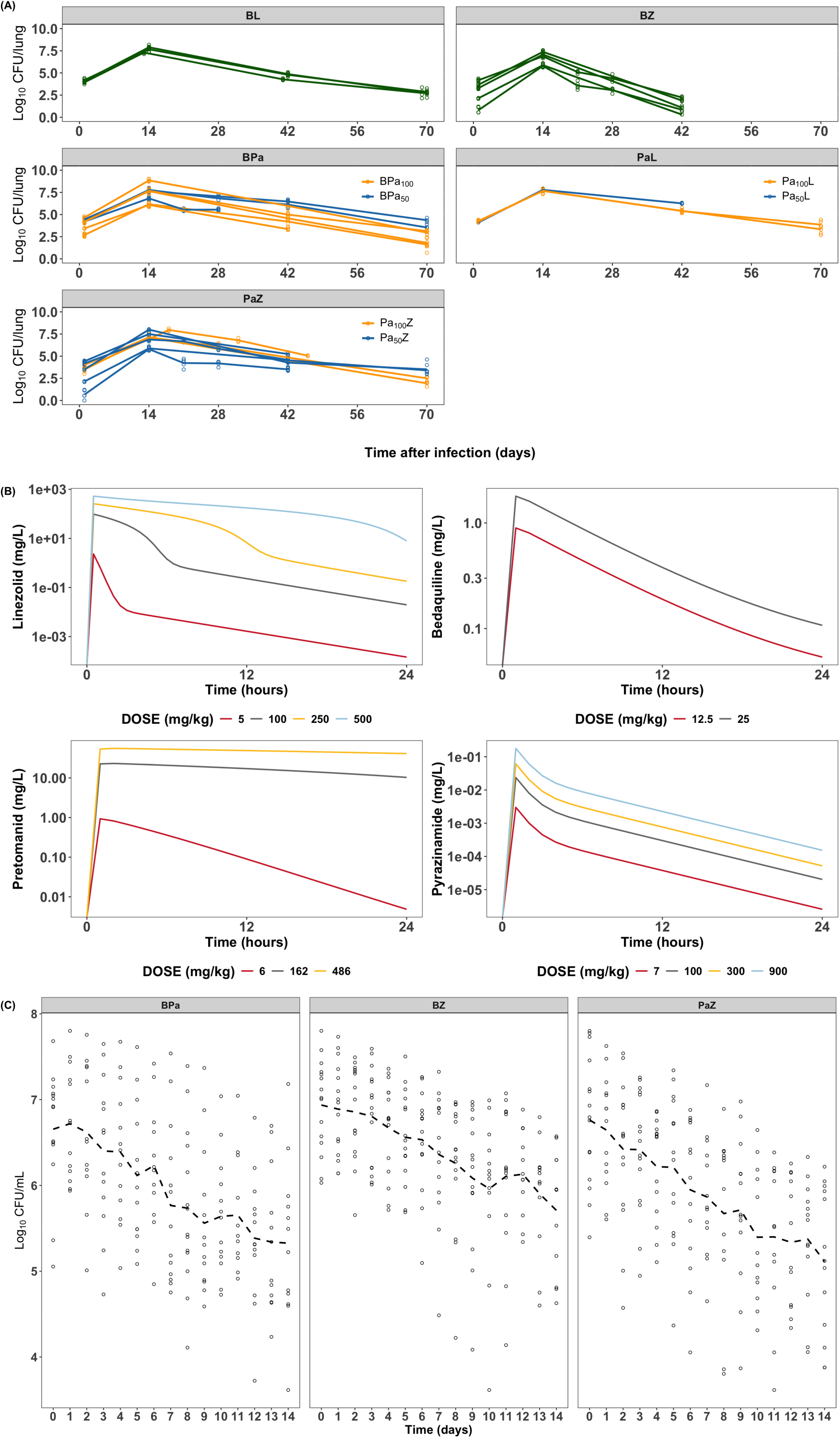
Observed data and model simulations used for model training and validation. Information on all doses is provided in Tables 1, 2, and 3. (A) Observed lung CFU counts in a high-dose aerosol-infection model in BALB/c mice inoculated with the drug-susceptible M. tuberculosis H37Rv strain on Day 0 and treated beginning on Day 14 with either BPa, BL, BZ, PaL, or PaZ. Pretomanid doses were 50 or 100 mg/kg, while other drugs were administered at fixed doses: bedaquiline 25 mg/kg, linezolid 100 mg/kg, and pyrazinamide 150 mg/kg. (B) Mouse PK simulations used to estimate the exposure-response relationship of the add-on SUPER drug to the backbone regimen. (C) Human Phase IIa early bactericidal activity study data for the BPa, BZ, and PaZ combinations. The dots represent observed values, and the dashed line represents the mean CFU decline over time across all patients from study NC-001. All doses were administered once daily.

**Table 1.** Mouse PK data used to estimate the PK-PD parameters in monotherapy regimens.

| Regimen | Doses (mg/kg) |
| --- | --- |
| **B** | 12.5, 25 single dose |
| **Pa** | 6, 162, 486 single dose;<br>100 daily for 4 or 8 weeks |
| **L** | 5, 100, 250, 500 single dose |
| **Z** | 7, 100, 900 single dose |

\*Reference (10)

**Table 2.**
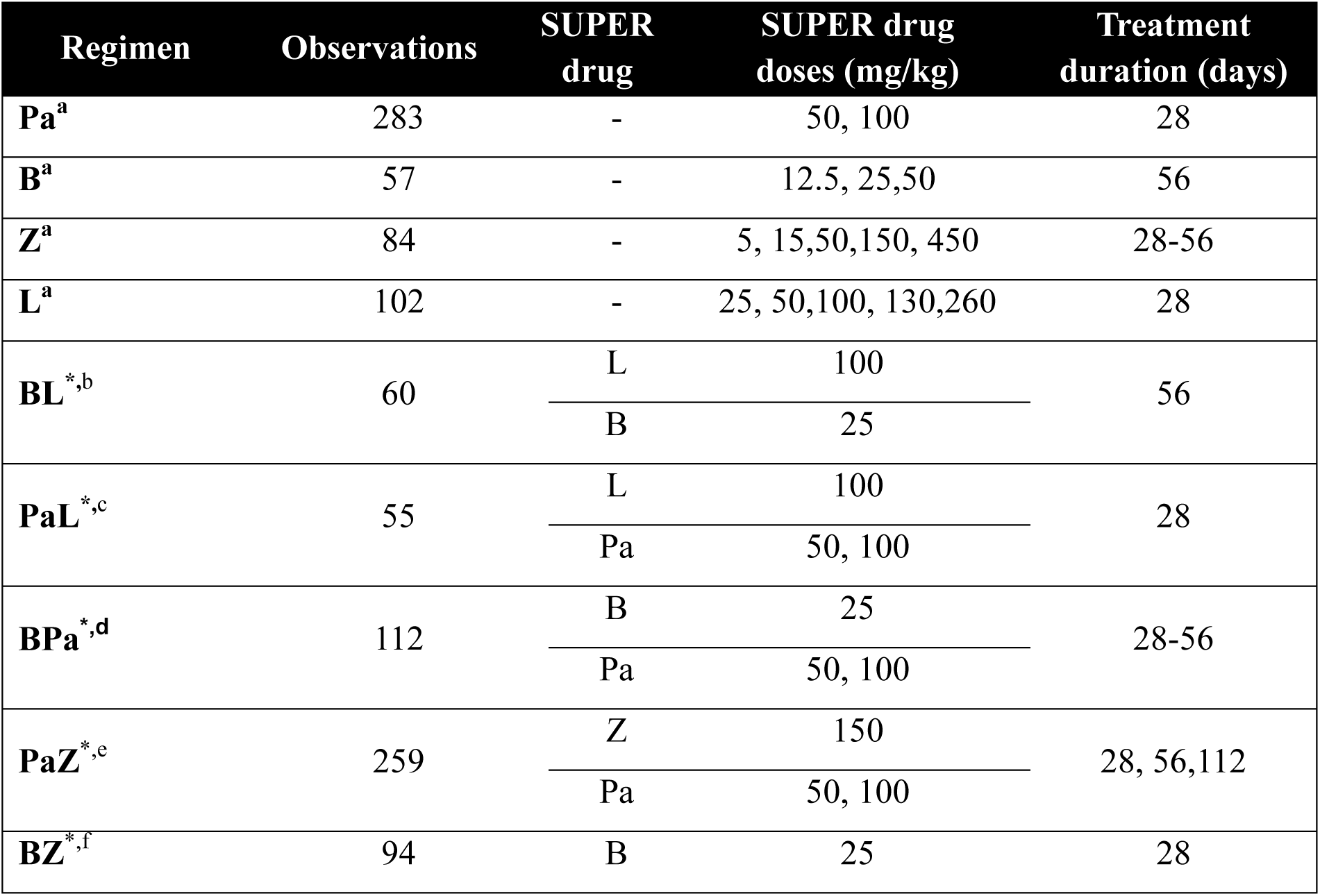
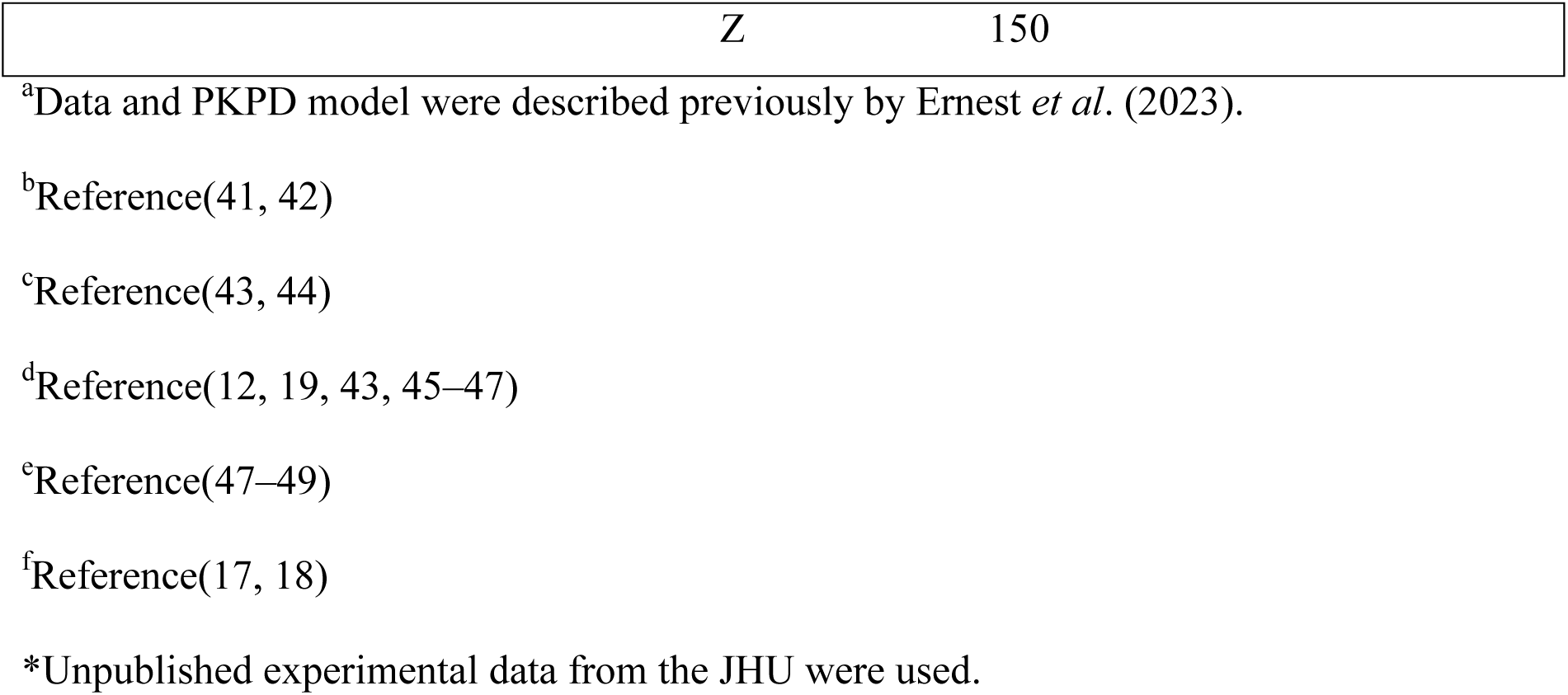
Mouse PD data.

| Regimen | Observations | SUPER drug | SUPER drug doses (mg/kg) | Treatment duration (days) |
| --- | --- | --- | --- | --- |
| **Pa<sup>a</sup>** | 283 | - | 50, 100 | 28 |
| **B<sup>a</sup>** | 57 | - | 12.5, 25, 50 | 56 |
| **Z<sup>a</sup>** | 84 | - | 5, 15, 50, 150, 450 | 28-56 |
| **L<sup>a</sup>** | 102 | - | 25, 50, 100, 130, 260 | 28 |
| **BL<sup>\*,b</sup>** | 60 | L | 100 | 56 |
|  |  | B | 25 |  |
| **PaL<sup>\*,c</sup>** | 55 | L | 100 | 28 |
|  |  | Pa | 50, 100 |  |
| **BPa<sup>\*,d</sup>** | 112 | B | 25 | 28-56 |
|  |  | Pa | 50, 100 |  |
| **PaZ<sup>\*,e</sup>** | 259 | Z | 150 | 28, 56, 112 |
|  |  | Pa | 50, 100 |  |
| **BZ<sup>\*,f</sup>** | 94 | B | 25 | 28 |

|  |  |
| --- | --- |
| Z | 150 |

<sup>a</sup>Data and PKPD model were described previously by Ernest *et al.* (2023).
<sup>b</sup>Reference(41, 42)
<sup>c</sup>Reference(43, 44)
<sup>d</sup>Reference(12, 19, 43, 45–47)
<sup>e</sup>Reference(47–49)
<sup>f</sup>Reference(17, 18)
\*Unpublished experimental data from the JHU were used.

PK simulations in mice were performed to estimate the exposure-response relationship of the drug of interest (i.e., add-on SUPER drug) to the backbone regimen when the SUPER approach was implemented in the analysis (Fig. 2B). To validate the translational PK-PD models for prediction of clinical trial outcomes, human sputum CFU counts over 14 days of treatment were collected from the Phase IIa NC-001 study (PaZ, BPa, and BZ) (Table 3, Fig. 2C) (16).

**Table 3.**
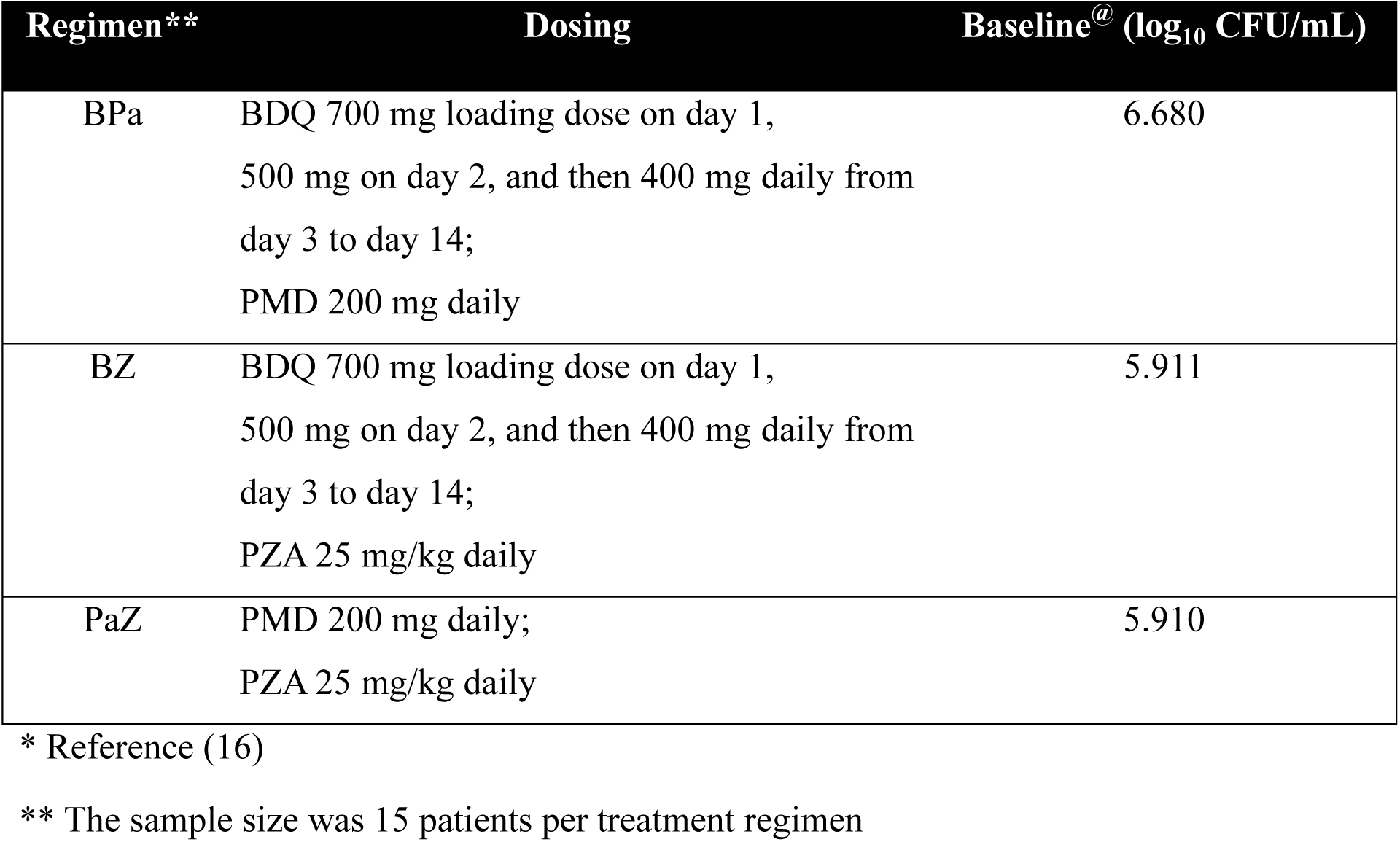
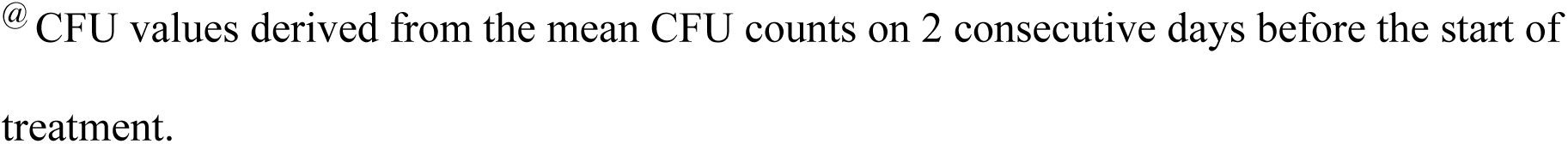
Clinical PD observations from the NC-001 trial (NCT01215851) used to validate the model*.

| Regimen** | Dosing | Baseline<sup>@</sup> (log<sub>10</sub> CFU/mL) |
| --- | --- | --- |
| BPa | BDQ 700 mg loading dose on day 1,<br>500 mg on day 2, and then 400 mg daily from<br>day 3 to day 14;<br>PMD 200 mg daily | 6.680 |
| BZ | BDQ 700 mg loading dose on day 1,<br>500 mg on day 2, and then 400 mg daily from<br>day 3 to day 14;<br>PZA 25 mg/kg daily | 5.911 |
| PaZ | PMD 200 mg daily;<br>PZA 25 mg/kg daily | 5.910 |

\* Reference (16)
\*\* The sample size was 15 patients per treatment regimen
@ CFU values derived from the mean CFU counts on 2 consecutive days before the start of treatment.

### Modeling of pharmacodynamic interactions

Depending on the amount of available data, either the GPDI or SUPER approach was used to quantify PD interactions between drugs in two-drug combinations. All models were evaluated using goodness-of-fit plots and prediction-corrected visual predictive check plots for both no-treatment and combination therapy conditions (Supplementary Fig. S1 and S2). The results showed that the observed data consistently fell within the 95% prediction interval of bacterial counts in the final PK-PD models for both approaches.

### Empirical method, SUPER approach, linked to the PK-PD translation platform

The SUPER approach (7) allowed us to characterize the effect of adding a companion drug to the drug of interest (the SUPER drug). Several models were evaluated to assess the impact of the companion drug on the E_max_ and/or EC_50_ of the SUPER drug. As a starting point, a simpler model was evaluated, assuming a homogeneous bacterial population and estimating a single value for EC_50_ or E_max_. Additionally, a mechanistic model was used to estimate the changes in the PD parameters of the SUPER drug in both the fast-replicating bacterial sub-population (during the first 28 days of treatment) and the slow-replicating bacterial sub-population (after 28 days).

The PK–PD model that best described the BL data, assuming linezolid as the SUPER drug, considered the drug effect as inhibitory on bacterial growth (Equation 2), whereas for the other tested combinations, the drug effect was better captured as an induction of bacterial death (Equation 3). Combinations in which different doses of the companion drug were evaluated were treated as independent regimens. Each drug was considered as the SUPER drug, allowing for an analysis of how its efficacy changes following the addition of the companion drug (Fig. 3).

**Figure 3.**
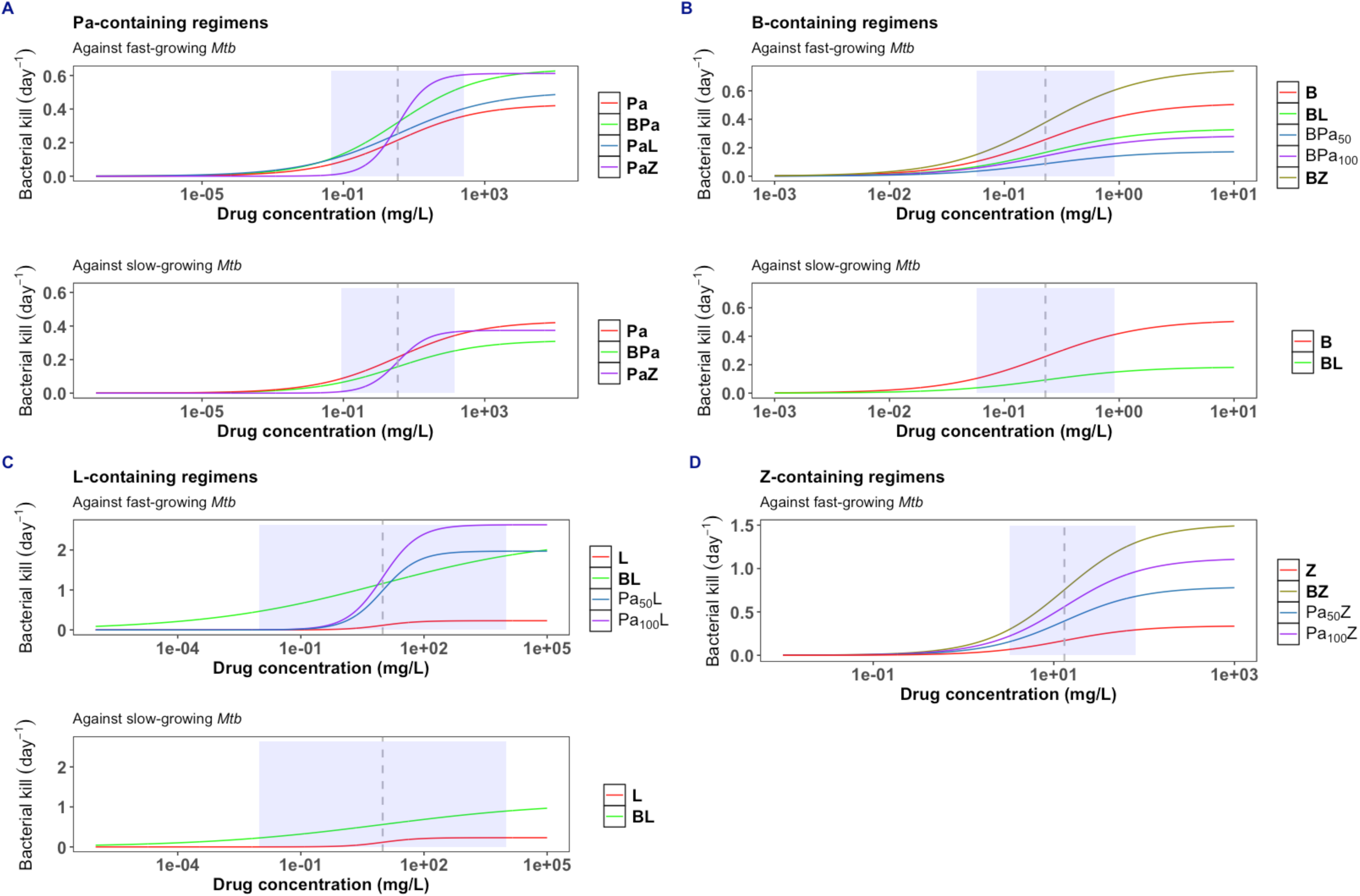
Exposure-response relationships in mice for each drug (bedaquiline, pretomanid, linezolid, and pyrazinamide) modeled as the SUPER drug in different two-drug combinations using the SUPER approach. Each panel represents a distinct two-drug combination: (A) pretomanid-based, (B) bedaquiline-based, (C) linezolid-based, and (D) pyrazinamide-based. The shaded area represents the simulated range of mouse plasma concentrations producing between 20% and 80% of the maximal effect, used to characterize the exposure–response relationship of the add-on SUPER drug to the backbone drug.

The model that best described the decline in log_10_ CFU profiles in mice was the E_max_ model, in which the EC_50_ of the SUPER drug was fixed at its monotherapy-estimated value, and only the change in E_max_ for the combination was estimated. Whenever possible, the change in E_max_ for the SUPER drug was estimated for both fast-replicating and slow-replicating *Mtb* sub-populations, represented as E_max,fast_ and E_max,slow_, respectively. When this was not possible owing to limitations in the availability of existing data for a given combination, a single E_max_ value was estimated. The parameter estimates and associated relative standard errors (RSEs) are shown in Table 4.

**Table 4.**
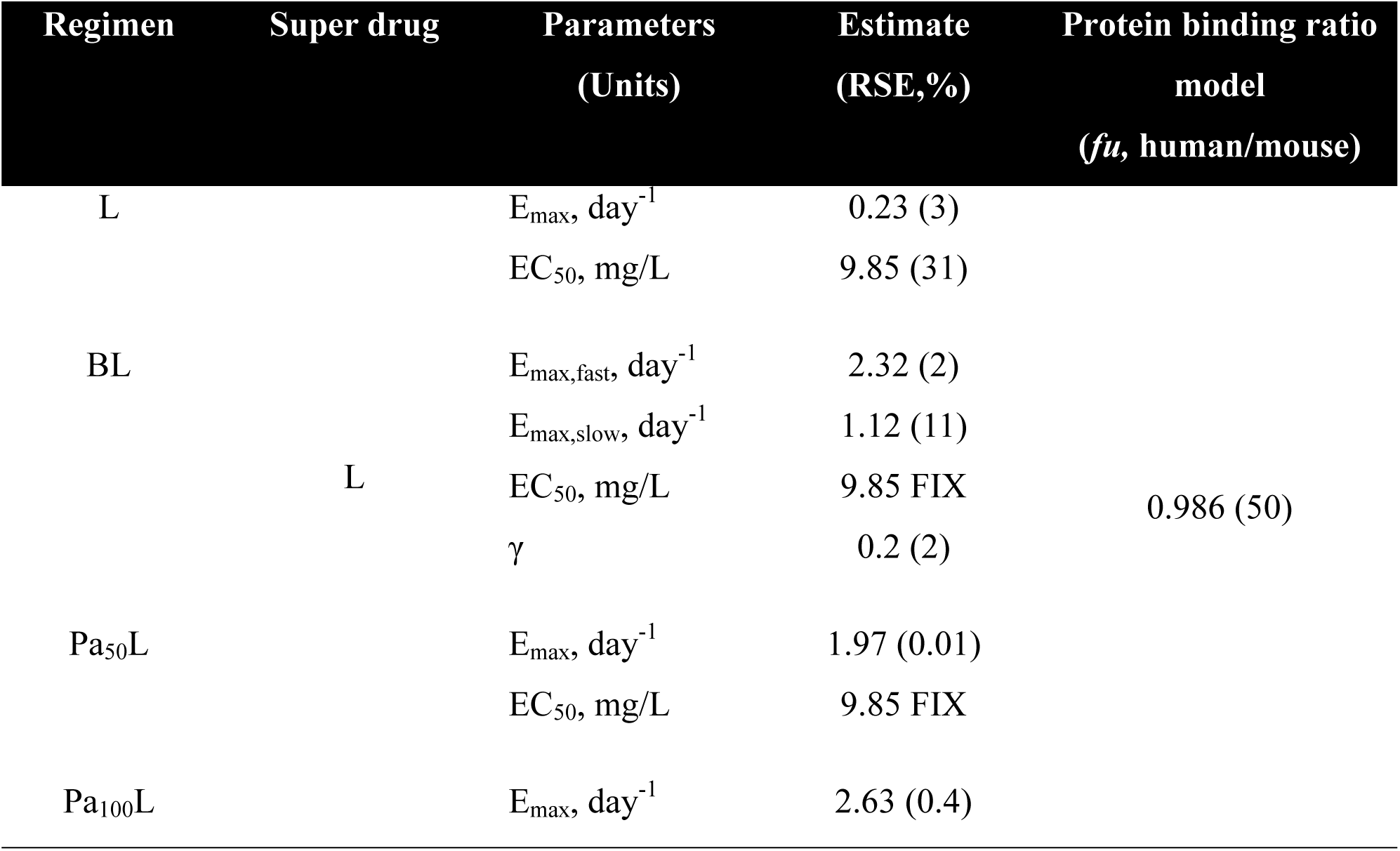
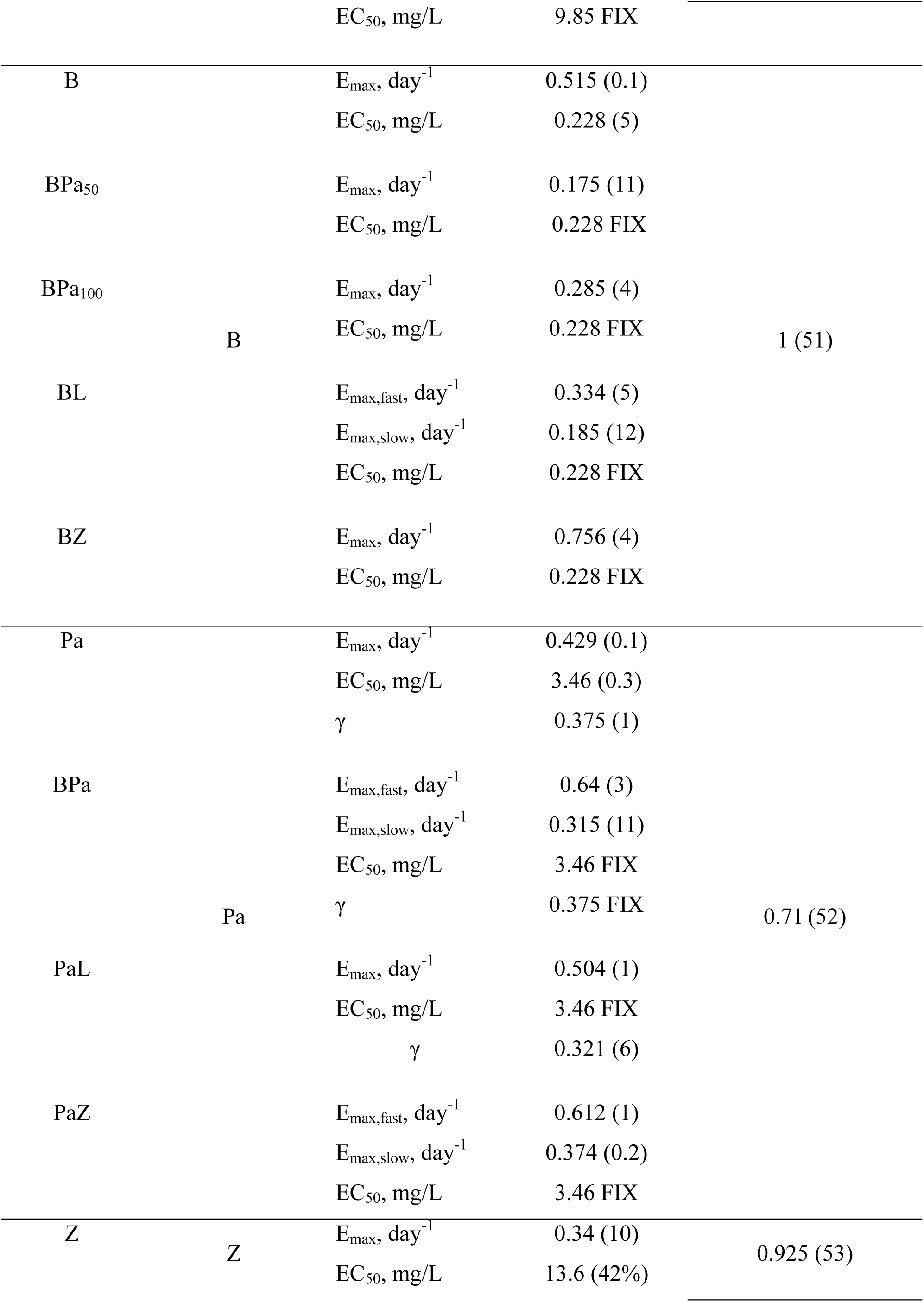
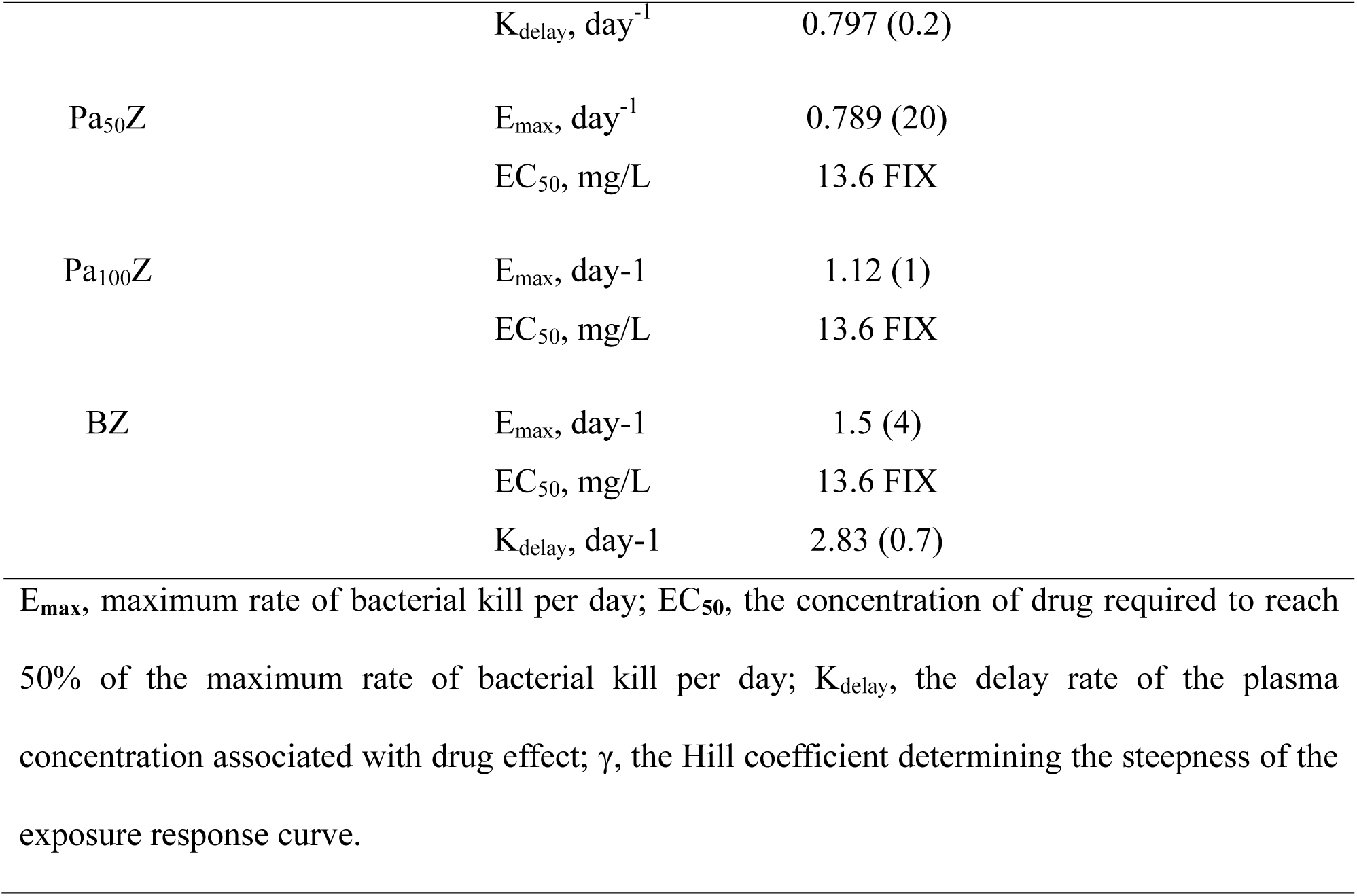
Mouse PK-PD model parameters using monotherapy and combinations.

For pretomanid-containing combinations, specifically for BPa and PaZ, the model differentiating E_max,fast_ and E_max,slow_ provided a better fit to the log_10_ CFU profiles in mice than the simpler model. The addition of pyrazinamide (PaZ) or bedaquiline (BPa) to pretomanid resulted in an approximately 1.4-fold increase in maximum efficacy during the first 28 days of treatment, ranging from 0.429 day^-1^ to 0.64 day^-1^ and 0.612 day^-1^, respectively. However, a decrease in maximum efficacy was observed after 28 days of treatment (Table 4). In contrast, the addition of linezolid (PaL) had little effect on pretomanid activity, with only a minor improvement observed (Fig. 3A).

While most two-drug combinations led to an increase in maximum efficacy, this effect was not observed when bedaquiline was paired with pretomanid or linezolid, according to our translational PK-PD model (Fig. 3B). Regarding bedaquiline-containing combinations, the additive effect of pyrazinamide (BZ) showed it to be the most effective companion drug for bedaquiline. The addition of linezolid (BL) or pretomanid to bedaquiline (BPa), reduced the maximum efficacy estimates (Fig. 3B).

The maximum efficacy of linezolid increased significantly with the addition of bedaquiline or pretomanid. The effect of adding pretomanid to linezolid (PaL) could not be evaluated for the slowly replicating bacterial sub-population due to a lack of data beyond 28 days of treatment. It was observed that combining pretomanid and linezolid improved the antibacterial activity of linezolid in a dose-dependent manner (9- to 11-fold). The addition of bedaquiline to linezolid (BL) led to an increase in E_max_ for both sub-populations, with changes of 10-fold in E_max,fast_ and 5-fold in E_max,slow_ (Fig. 3C).

For pyrazinamide-containing combinations, the simpler model, which assumed a homogeneous bacterial population with a single estimated E_max_ value, provided a better fit to the log_10_ CFU profile in mice than a model estimating E_max,fast_ and E_max,slow_. The addition of pyrazinamide to pretomanid (PaZ) or bedaquiline (BZ) enhanced the antibacterial activity of pyrazinamide by nearly 2- and 4-fold, respectively (Fig. 3D). As observed for the addition of pretomanid in the PaL combination, the effect of pretomanid when added to bedaquiline or pyrazinamide followed a similarly dose-dependent pattern.

Overall, it was observed that in the two-drug combination where the E_max_ value could be estimated for each bacterial sub-population, the addition of a companion drug had a more pronounced effect on the E_max_ of the SUPER drug in the fast-replicating bacterial sub-population compared to the slow-replicating one (Fig. 3). In summary, these results suggested that pyrazinamide combined with pretomanid or bedaquiline constitutes a highly potent backbone for novel regimens. The changes in the maximum rate of bacterial kill per day (E_max_, day⁻¹) estimates for each drug when administered in different two-drug combinations using the SUPER approach are shown in Figure S3 (Supplementary Material).

### General Pharmacodynamic Interaction (GPDI) model linked to the PK-PD translation platform

To assess and describe the PD interactions between bedaquiline, pretomanid, linezolid, and pyrazinamide in two-drug combinations, the translational PK-PD model was linked to the GPDI model. PD interactions were evaluated by quantifying the fractional changes in either EC_50_ or E_max_. The final parameter estimates for each drug combination are summarized in Table 5.

**Table 5.**
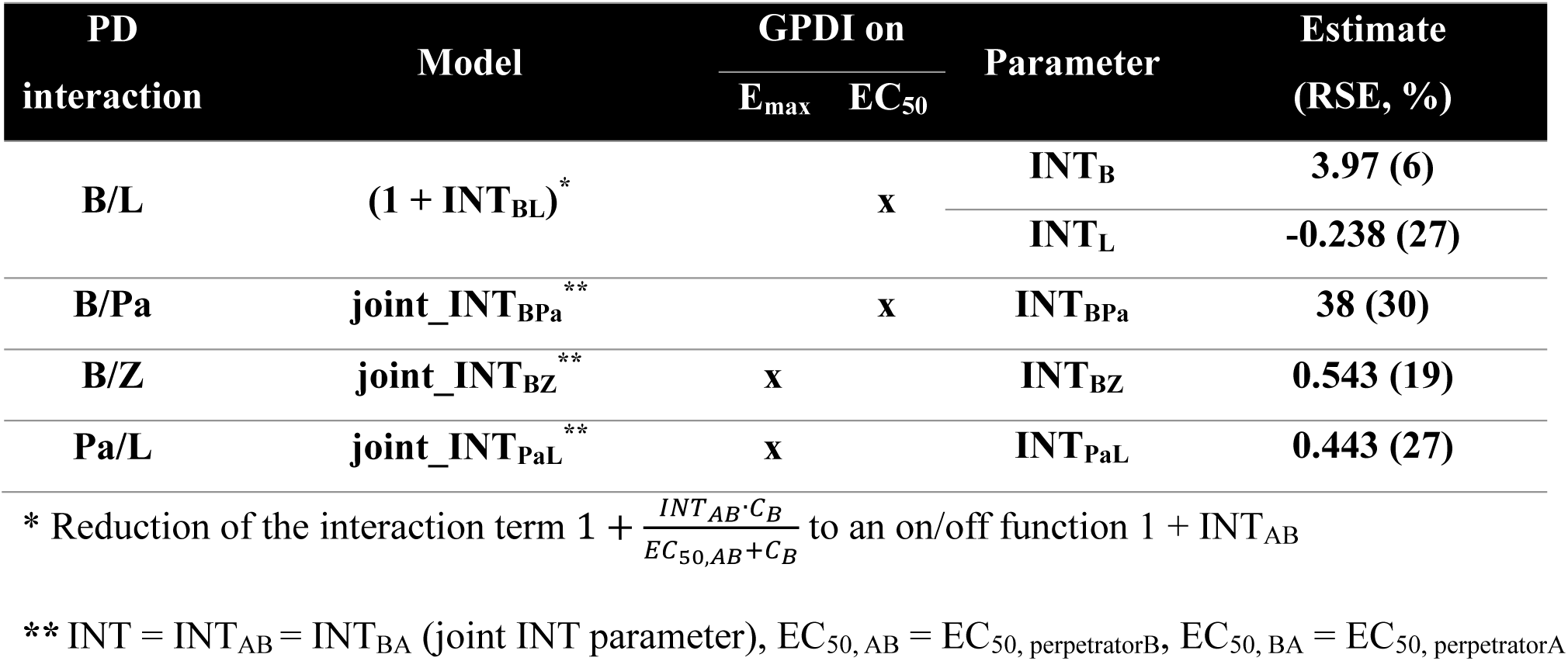
Parameter estimates of final interaction model.

Using the simplified on/off GPDI interaction function (1 + INT_AB_) implemented on EC_50_, asymmetric interactions were identified in the BL combination. However, for the two-drug combinations BPa, BZ, and PaL, a joint interaction parameter (joint_INT) was estimated to describe drug interactions. This modeling assumption simplified the model by assuming bidirectional interactions, which prevented capturing asymmetric interactions.

Bedaquiline exhibited a decrease in potency when combined with linezolid (INT_B_ = 3.97), indicating an approximately 5-fold increase in EC_50_ of bedaquiline in the presence of linezolid. This suggests that when combined with linezolid, bedaquiline required higher exposure to achieve the same level of bacterial killing as in monotherapy. Conversely, linezolid displayed a modest increase in potency when combined with bedaquiline, with an INT_L_ estimate of −0.238, indicating a shift toward lower EC_50_ values and thus enhanced potency.

Figure 4 shows simulations of the log₁₀ CFU/lung at 28 days and the PD interaction classification of this biomarker from the final translational PK-PD-GPDI model. For the BL combination, antagonism persisted across bedaquiline doses and showed a modest reduction at the highest linezolid dose. These PD interactions resulted in up to 1.65 log_10_ CFU/lung higher than the expected additive effect, reflecting moderate antagonism in BL combinations (Fig. 4A). Antagonism was observed for all BPa combinations and was more pronounced at higher pretomanid doses, suggesting concentration-dependent interactions (Fig. 4B).

**Figure 4.**
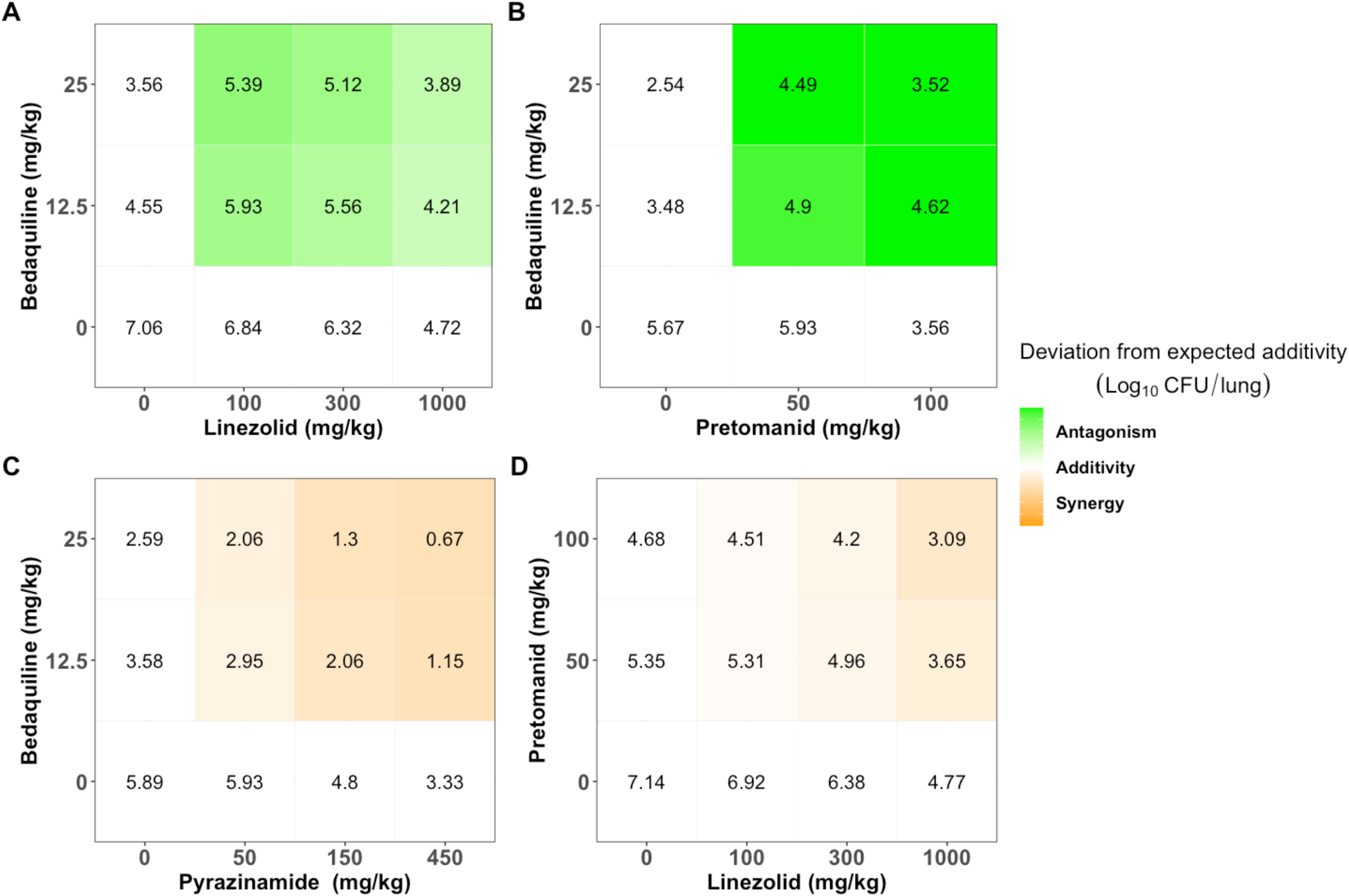
Predicted log_10_ CFU/lung in numbers and log_10_ CFU/lung deviation from expected additivity (in shaded areas) for two-drug combinations: bedaquiline and linezolid (A), bedaquiline and pretomanid (B), pyrazinamide and bedaquiline (C), and pretomanid and linezolid (D) at 28 days after treatment. White areas in the figure show expected additivity, whereas green shaded areas show higher log_10_ CFU/lung than expected additivity (antagonism), and orange shaded areas show lower log_10_ CFU/lung than expected additivity (synergism).

Bedaquiline and pyrazinamide increased their E_max_ when combined (Fig. 4C), as reflected by a joint_INT_BZ_ estimate of 0.543 using the GPDI model. Similarly, pretomanid and linezolid demonstrated an enhanced antibacterial effect when combined (Fig. 4D), with a joint_INT_PaL_ estimate of 0.443. These results indicate that both BZ and PaL achieved greater bacterial killing compared to the effects observed with each drug alone. The data did not support any PD interaction between pretomanid and pyrazinamide. The VPCs for no treatment and combination therapies based on the final models are shown in Figure S2.

### Translational simulations for prediction of Phase IIa outcomes

Using exposure-response relationships derived from the mouse model and correcting for differences in the fraction of unbound drug in plasma between mice and humans for EC_50_ estimates, the translational model predicted the clinical 14-day sputum CFU count in NC-001 for the combinations BPa, PaZ, and BZ with reasonable accuracy using the SUPER approach. In general, the model prediction intervals fit well with the observed clinical data, with the median prediction closely aligned with the median observed log_10_ CFU/mL decline for up to 7 days of treatment (Fig. 5A). However, beyond 7 days, a slight underprediction was observed for PaZ and BZ (with Z as the SUPER drug), and BPa (with Pa as the SUPER drug), while the model tended to overpredict CFU decline for BZ (with B as SUPER drug). For most combinations, predictions remained within the 95% prediction interval throughout the treatment period, although predictions for BZ (with B as the SUPER drug) fell outside this interval toward the end of treatment.

**Figure 5.**
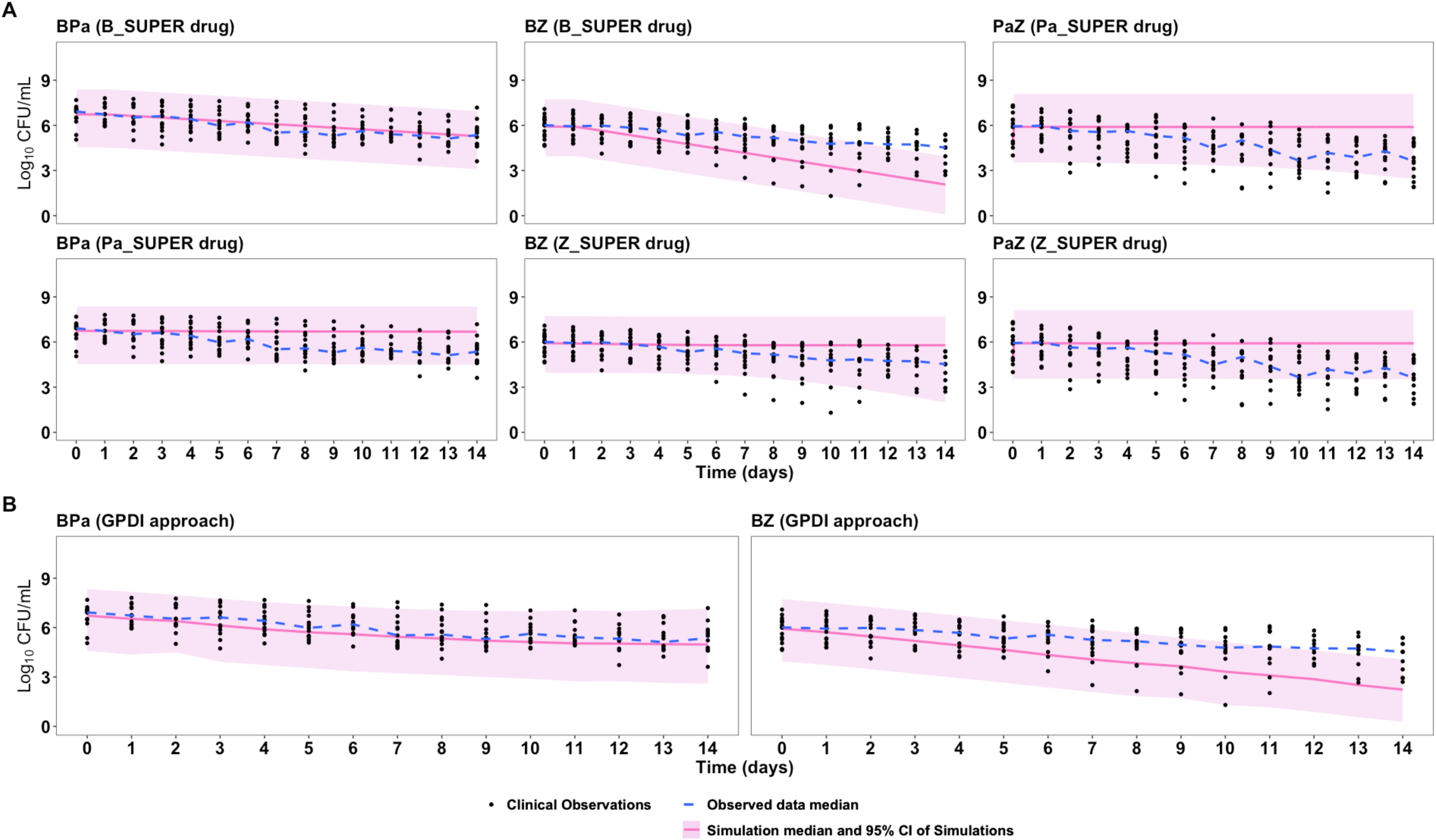
Phase IIa predictions at 14 days of treatment. (A) Predicted early bactericidal activity of bedaquiline and pretomanid, pyrazinamide and bedaquiline, and pyrazinamide and pretomanid combinations in the NC-001 trial by PK-PD relationships determined in mice using only the SUPER approach. (B) Predicted early bactericidal activity of bedaquiline and pretomanid and pyrazinamide and bedaquiline combinations in the NC-001 trial by PK-PD relationships determined in mice using only the GPDI approach. Solid line represents median of model prediction, and dashed line represents median sputum count of clinical data.

These reasonably accurate predictions, achieved without individually modeling the effects of each drug, highlight the capability of the SUPER approach to reliably characterize the exposure-response of drug combinations for further application. For the BZ combination, the model tended to overpredict the log₁₀ CFU/mL decline. When the GPDI approach was applied, similar results were observed for the BPa and BZ combinations (Fig. 5B, Fig. S4).

## Discussion

A comprehensive understanding and precise characterization of PD interactions is crucial for the rational selection of drug combinations during the early stages of drug development. In TB, where multidrug regimens are essential, the ability to accurately predict drug interactions can significantly impact treatment efficacy, duration, and success. This study addresses this need by evaluating two complementary modeling approaches, the empirical SUPER method (7) and the mechanistic GPDI model (13), to characterize drug interactions in TB combination therapies.

Using BPaMZ as a case study (11), the predictive capacity of our translational PK-PD platform for drug combinations, originally developed to predict short-term monotherapy clinical outcomes, was previously validated through integration of the SUPER method. It was demonstrated that this platform could predict Phase IIb sputum CFU counts over 8 weeks as well as Phase III outcomes.

In this study, we extended this validation by demonstrating that our translational PK-PD platform can also predict clinical outcomes in Phase IIa trials of drug combinations against TB. Importantly, we implemented two distinct approaches by separately linking either the empirical SUPER method or the mechanistic GPDI model to our translational PK-PD platform. Each approach was implemented independently, allowing us to evaluate their individual predictive performance. Preclinical murine model data for drug combinations were used to train PK-PD models, capturing the combination efficacy of the regimens and successfully predicting clinical EBA for BZ, PaZ, and BPa using each approach separately.

Our analyses revealed important findings regarding two-drug combinations. Across both independent models, consistent identification of antagonistic interactions emerged when bedaquiline was combined with either pretomanid or linezolid. Specifically, a reduction in bedaquiline activity was detected, manifested as decreased potency through the GPDI model and reduced E_max_ with the SUPER method when combined with either pretomanid or linezolid.

Asymmetric interactions were observed for the BL combination, even though a single dose level of each drug was tested. While the potency of bedaquiline declined with the addition of linezolid, linezolid showed a modest increase in potency in the presence of bedaquiline. This was particularly evident in the GPDI estimates (INT parameter), and the SUPER model captured the enhanced efficacy parameters for linezolid. For the two-drug combinations BPa, BZ, and PaL, a joint interaction parameter (joint_INT) was estimated to describe the drug interactions. This modeling assumption incorporated bidirectional interactions, thereby restricting the capacity of the model to accurately represent asymmetric interaction dynamics. Both drugs influenced each other to the same extent and in the same direction. Considering that some drug combinations have been shown to exhibit asymmetric interactions in vivo, this simplification represents a limitation of this modeling approach. The antagonism of pretomanid on bedaquiline activity observed in our analysis aligns with previous experimental findings (17–22). Tasneen *et al*. (17–19) reported that, in BALB/c mice infected with *Mtb* H37Rv, the combination of bedaquiline and pretomanid exhibited lower bactericidal activity than bedaquiline alone during the first month of treatment. Similar observations were made for the *Mtb* Erdman (23) and HN878 (21) strains in this model.

Simulations from the final translational PK-PD GPDI model showed an antagonistic interaction between bedaquiline and both linezolid and pretomanid. BL antagonism was consistent across all simulated doses, whereas BPa antagonism varied with drug exposure levels. At higher pretomanid doses, increasing bedaquiline doses reduced antagonism, with the most favorable interaction observed at the highest doses of both drugs. This dose-dependency suggests that careful optimization of dosing strategies could mitigate negative interactions and preserve the therapeutic efficacy.

Using both modeling approaches, we identified pyrazinamide as the most effective companion drug for bedaquiline and pretomanid. The SUPER approach demonstrated that pyrazinamide significantly enhanced the antibacterial activity of both drugs; however, changes in drug potency could not be estimated, only changes in E_max_. The synergism between pyrazinamide and bedaquiline observed in our analysis is further supported by studies in Swiss mice infected with *Mtb* H37Rv (24) and in both BALB/c and C3HeB/FeJ mouse infection models infected with *Mtb* Erdman (25). Despite differences in pulmonary pathology between these models, the BZ combination showed significant synergistic activity during the initial two months of treatment.

Similarly, the GPDI model showed that the BZ combination substantially increased maximum efficacy. These consistent results across different modeling methodologies underscore the potential of pyrazinamide as a key backbone agent in multidrug regimens. Goh *et al*. (11) demonstrated that the combination of pretomanid with pyrazinamide enhanced both potency and E_max_ compared to pretomanid monotherapy. Furthermore, Muliaditan *et al*. (7) reported similar effects for the potency of bedaquiline when combined with pyrazinamide for both fast- and slow-growing bacterial subpopulations, while the potency of bedaquiline decreased for both subpopulations when combined with pretomanid.

These findings are consistent with the clinical EBA reported by Diacon *et al*. (16). In this Phase IIa trial, the addition of pyrazinamide markedly improved the EBA of bedaquiline between days 2 and 14 (EBA_2-14_), increasing it from 0.076 to 0.143. Additionally, the EBA of the combination from day 7 to day 14 showed an important increase compared with bedaquiline monotherapy (0.123 to 0.152), even as the EBA of bedaquiline alone increased during the second week of treatment. In contrast, the addition of pretomanid did not increase the EBA_7-14_ of bedaquiline (0.123 to 0.114).

The SUPER and GPDI approaches, when individually linked to our translational PK-PD platform, offer complementary insights into drug interactions. The SUPER approach allowed us to quantify how the addition of a companion drug modified the efficacy of the SUPER drug. This approach is particularly valuable when limited data are available, or when a simplified model is sufficient for predictive purposes. Our results showed that the SUPER approach, when linked to the translational platform, accurately predicted clinical outcomes for various two-drug combinations, supporting its utility in early-phase development.

The GPDI model, built on a mechanistic basis, was independently integrated into the same translational platform to provide an understanding of the nature and directionality of drug interactions. By quantifying how drugs alter each other’s potency or efficacy, it enables analysis of asymmetric interactions and dose-dependent effects. PD interactions were evaluated using the GPDI model under the Bliss Independence (BI) criterion (26), selected due to the distinct mechanisms of action of the drugs, their potential for independent activity, and differences in their maximum achievable efficacy.

When comparing the SUPER and GPDI approaches integrated into our translational PK-PD platform, we observed that each method offers distinct advantages for characterizing drug interactions. The SUPER approach provides a practical framework that requires fewer parameters and less extensive data, making it particularly suitable for early-stage drug development when data may be limited. This approach excelled at identifying how companion drugs modified the efficacy of the drug of interest and provided important insights into differential effects on bacterial subpopulations.

Although our models demonstrated reasonable precision in predicting Phase IIa outcomes, evidenced by prediction intervals fitting well with observed clinical data and median predictions closely aligned with the median observed log_10_ CFU/mL decline, they tended to overpredict the EBA for the BZ combination when bedaquiline was assumed to be the SUPER drug. This tendency to overpredict the effect of bedaquiline has been previously reported, both in Phase IIa predictions for bedaquiline monotherapy (10) and in Phase IIb simulations of B-containing combinations such as BPaZ and BPaMZ (11). One possible explanation, suggested by earlier studies, is that the active metabolite of bedaquiline, M2, was not incorporated into the models. In mice, bedaquiline is extensively metabolized to M2, which likely plays a significant role in the bactericidal effects. Since both compounds have shown additive bactericidal effects in vitro and in vivo (28), both bedaquiline and M2 contribute substantially to bactericidal activity in mice. Conversely, M2 formation is less extensive in humans, resulting in minimal contribution to overall efficacy (27, 28). The interspecies differences in the BDQ to M2 metabolite ratio between mice and humans are likely relevant for efficacy prediction in translational models (10). This limitation could be addressed in the PK-PD models developed in this study by explicitly incorporating active metabolites and accounting for interspecies PK differences.

However, no overprediction was observed for BPa, suggesting that other factors could explain the findings observed for BZ. The observed overprediction of BZ efficacy when translating from murine models to humans may be explained by interspecies differences in lesion pathology and microenvironmental conditions, as supported by previous experimental evidence (25, 29, 30). TB patients present a wide spectrum of morphologically distinct pulmonary lesion types, ranging from cellular to caseous, necrotic, and hypoxic granulomas, which create heterogeneous microenvironments that influence drug efficacy (26). In contrast, BALB/c mice develop uniform, non-necrotic lesions that do not replicate the complex microenvironment characteristic of human disease. Evidence from studies in BALB/c and C3HeB/FeJ mice, the latter developing caseous and necrotic granulomas more representative of human TB, has shown variable responses to BZ and pyrazinamide monotherapy (25, 29, 30). The BZ combination produced a potent and uniform response in BALB/c mice; however, in C3HeB/FeJ mice, it resulted in a bimodal response with subgroups of lesions or animals showing reduced sensitivity, reflecting the variability observed in patients. These findings have been linked to microenvironmental factors such as neutral pH, hypoxia, and altered bacterial metabolism, all of which diminish pyrazinamide activity despite adequate tissue penetration. Additionally, bedaquiline and its M2 metabolite showed lower and slower accumulation in necrotic caseous tissue compared with cellular lesions (26). Taken together, these data suggest that the simplified and homogeneous lesion microenvironment in BALB/c murine models may contribute to an overestimation of the efficacy of the BZ combination when extrapolated to the heterogeneous pathology of human tuberculosis.

Our simulated EBA for the BPa combination closely matches the EBA observed in the NC-001 trial. The addition of pretomanid did not increase (and may have decreased) the EBA_7-14_ of bedaquiline (0.123 - 0.114), consistent with the predicted antagonistic interaction (16). Indeed, the EBA of pretomanid alone at 200 mg/day, previously shown to be 0.106 and 0.109 in trials performed at the same trial site, may have limited any further negative effects of an antagonistic interaction (31). Moreover, the higher EBA of pretomanid (0.106 - 0.109) observed in the earlier trials compared to the EBA_0-7_ of bedaquiline alone (0.043) in NC-001 suggests that pretomanid primarily drives the observed EBA over the first week of BPa treatment (EBA_0-7_ of 0.114 in NC-001), possibly with a small additive effect of bedaquiline. Meanwhile, the EBA of bedaquiline increases over time as its exposure at the site of action rises. Thus, the contribution of individual drugs to the BPa combination appeared to change over the first 14 days in a manner that was consistent with their predicted interaction.

Limited clinical data for two-drug combinations, especially from Phase IIa studies, prevented validation of the BL and PaL translational models. Moreover, it was not possible to simulate the EBA for the PaZ combination using the GPDI approach because the data did not support a reliable estimation of PD interactions for this combination.

Considering the limitations identified in this study for the GPDI model, we suggest using an alternative translational approach that combines our PK-PD platform (10) with the SUPER approach. This integrated framework performed well in predicting clinical Phase IIa outcomes even with limited preclinical data. We also showed that it can predict short-term outcomes for combinations, making it a practical tool to guide decisions throughout different stages of drug development.

## Methods

### Data Collection for PK-PD Model Development and Validation

The general workflow of modeling and simulation is shown in Figure 1. A detailed list of preclinical data included in the current analysis can be found in Tables 1 and 2. PK data from mice treated with bedaquiline (B), pretomanid (Pa), pyrazinamide (Z), and linezolid (L) as monotherapy were collected from literature (10). Lung colony-forming unit (CFU) counts in mice after treatment were collected as efficacy data for both monotherapy and five two-drug combinations (BL, PaL, BPa, PaZ, BZ), as well as for natural bacterial growth (i.e., no treatment). Experiments were conducted in BALB/c mice infected with *Mtb* H37Rv strain using a subacute high-dose aerosol infection model to assess all drug combinations. The human population PK data for pyrazinamide (32), bedaquiline (33), pretomanid (34), and linezolid (35) were simulated based on published clinical PK models. Sputum CFU counts for PaZ, BPa, and BZ in a Phase IIa trial were collected from a published clinical study (16) with individual CFU data from NC-001 obtained from the TB Alliance.

### Model evaluation and selection

All model development and simulations were conducted using NONMEM (version 7.5) and Perl speaks NONMEM (PsN, 5.3.1) implementing first-order conditional estimation with interaction (FOCE-I) as the estimation method (36). An additive error model on a log scale was used to describe residual unexplained variability in the PD data. Model evaluation and selection were based on the objective function value (OFV), with a decrease of 3.84 considered statistically significant (P < 0.05, *χ*² distribution) for nested models with one degree of freedom. The Akaike information criterion (AIC) was used for the selection of non-nested models (37). Additionally, goodness-of-fit (GOF) plots, parameter precision, predictive performance assessed through visual predictive checks (VPC), prediction-corrected visual predictive checks (pcVPC) (38), and scientific plausibility, were used for model selection. VPCs and pcVPCs were generated using 1,000 model-based replicates, with the 5th, median, and 95th percentiles compared with the corresponding observed data to assess model performance. Data transformation and graphical outputs were performed in R (version 4.3.1) via the RStudio (2023.6.1) interface, applying the xpose4, ggplot, and tidyverse packages. The run record was generated with Pirana PMX Modeling (version 2.9.9).

### Modeling of pharmacodynamic interactions

To account for pharmacokinetic drug-drug interactions that could alter drug exposure, we ran all four drugs (bedaquiline, pretomanid, pyrazinamide, and linezolid) through the Drug Interactions Checker (39) and found no significant interactions. Therefore, we could safely assume that the pharmacokinetics of the drugs would not change significantly when used in two-drug combinations. Monotherapy mouse PK and PD were described using previously published integrated PK-PD models with bacterial dynamics from Ernest *et al*. for bedaquiline, pretomanid, pyrazinamide, and linezolid. (10) Similarly, as in our previously published translational work on monotherapy and the prediction of BPaMZ Phase IIb/III trial outcomes, mouse PK-PD models were developed by incorporating drug effects (EFF) into a previously published bacterial infection model that describes *Mtb* growth, death, and the adaptive immune response without drug treatment in BALB/c mice (Equation 1) (40). The bacterial killing rate (day⁻¹) was used to define the drug effect (EFF). Drug effects were tested both as an inhibitory effect on bacterial growth and as an induced bacterial death effect (Equations 2 and 3).

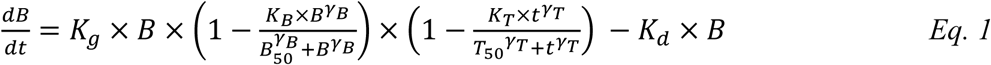

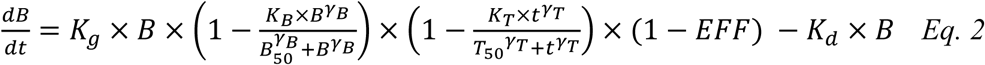

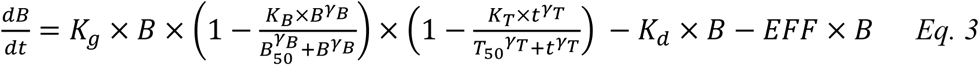

where *B* is the bacterial number; *K_g_* and *K_d_* characterize the bacterial growth rate and natural death rate, respectively; *K_B_*and *K_T_* represent the bacterial number-dependent maximal adaptive immune effect and incubation time-dependent maximal adaptive immune effect, respectively; *t* is the incubation time since inoculation; *B_50_*is the bacterial number that results in half of *K_B_*; *T_50_*is the bacterial number that results in half of *K_T_*; γ*_B_* and γ*_T_* are represent the steepness of the bacterial number-dependent and time-dependent immune effect relationships, respectively.

The effects of drug combinations were estimated using the general pharmacodynamics drug interaction (GPDI) model (13) and an empirical approach (SUPER) (7), based on the availability and abundance of data in the murine model. The drug effects of the combinations were evaluated in both fast- and slow-replicating bacteria. The full methodological details are provided in the Supplementary Material.

### Translational model for Phase IIa 14-day sputum

Mouse PK-PD relationships were translated to humans, and plasma protein binding ratios between humans and mice 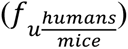 were used to convert unbound plasma drug concentrations from mice to humans, correcting for differences in protein binding between species (Table 5). Clinical PK simulations of bedaquiline, pretomanid, pyrazinamide, and linezolid (Table 3) were integrated with a translated exposure-response model from mouse PK-PD models to predict longitudinal changes in log₁₀ CFU over 14 days.

These simulations incorporated baseline demographics matching the Phase IIa NC-001 trial (16) patient population and accounted for the variability in patient PK profiles and baseline CFU. The net CFU change rate (K_net_) during the 14-day study was assumed to be zero, indicating that CFU changes were solely drug-driven (Equation 4) (10). Model predictions were evaluated by overlaying the mean and standard deviation lines from simulations with observed clinical data. Clinical outcomes were predicted by simulating CFU counts in TB patients using the translated PK-PD relationships, with 1000 simulations conducted for the BZ, PaZ, and BPa combinations, which had available clinical PD data.

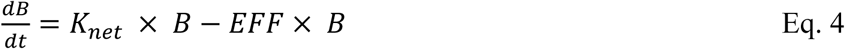

where K_net_ is the net rate of change in the bacterial number in the sputum of TB patients.

## Acknowledgements

We wish to thank the Tuberculosis Alliance (TBA) for providing the clinical data (NC-001 Phase IIa study). We would also like to thank Anu Patel, Ziran Li, Dongsheng Yang, and Samer Mouksassi for their feedback and discussions regarding this manuscript.

This work was supported by the National Institutes of Health/National Institute of Allergy and Infectious Diseases (grant number UM1 AI179699 “Preclinical Design and Clinical Translation of TB Regimens Consortium”, and grant number R01-AI-111992 “Integrative Pharmacology Approach to Understand Mechanisms of TB Drug Resistance project”); and by the TB Alliance (providing support for mouse model experiments conducted at Johns Hopkins University).

## Conflict of interest

Eric L. Nuermberger received research support (paid to Johns Hopkins University) from the TB Alliance. Eric L. Nuermberger served as a paid advisory board member for Johnson & Johnson.

## Author contributions

The author contributions were as follows:

Niurys de Castro Suarez, Formal analysis, Data curation, Investigation, Visualization, Writing – original draft | Jacqueline Ernest, Formal analysis, Visualization, Writing –review and editing | Eric L. Nuermberger, Investigation, Resources, Writing – review and editing | Rada Savic, Conceptualization, Funding acquisition, Methodology, Resources, Supervision, Writing – review and editing. All authors contributed to interpretation of the data and results, and read and approved the approved the final paper.

